# Genome-wide association analyses of multiple traits in Duroc pigs using low-coverage whole-genome sequencing strategy

**DOI:** 10.1101/754671

**Authors:** Ruifei Yang, Xiaoli Guo, Di Zhu, Cheng Bian, Yiqiang Zhao, Cheng Tan, Zhenfang Wu, Yuzhe Wang, Xiaoxiang Hu, Ning Li

## Abstract

High-density markers discovered in large size samples are essential for mapping complex traits at the gene-level resolution for agricultural livestock and crops. However, the unavailability of large reference panels and array designs for a target population of agricultural species limits the improvement of array-based genotype imputation. Recent studies showed very low coverage sequencing (LCS) of a large number of individuals is a cost-effective approach to discover variations in much greater detail in association studies. Here, we performed cohort-wide whole-genome sequencing at an average depth of 0.73× and identified more than 11.3 M SNPs. We also evaluated the data set and performed genome-wide association analysis (GWAS) in 2885 Duroc boars. We compared two different pipelines and selected a proper method (BaseVar/STITCH) for LCS analyses and determined that sequencing of 1000 individuals with 0.2× depth is enough for identifying SNPs with high accuracy in this population. Of the seven association signals derived from the genome-wide association analysis of the LCS variants, which were associated with four economic traits, we found two QTLs with narrow intervals were possibly responsible for the teat number and back fat thickness traits and identified 7 missense variants in a single sequencing step. This strategy (BaseVar/STITCH) is generally applicable to any populations and any species which have no suitable reference panels. These findings show that the LCS strategy is a proper approach for the construction of new genetic resources to facilitate genome-wide association studies, fine mapping of QTLs, and genomic selection, and implicate that it can be widely used for agricultural animal breeding in the future.

## Background

Genome-wide association studies (GWASs) have generated thousands of genetic variants associated with complex traits in human and agricultural species [1, 2]. The mapping resolution lies on the density of genetic markers which perceive linkage disequilibrium (LD) in sufficiently large populations [3, 4]. Several large-scale whole-genome sequencing projects have been completed, [5] which were designed to identify the underlying mechanisms that drive hereditary diseases in human and genomic selection in the breeding of agricultural species [6-8]. Despite the declining cost of sequencing, it is still difficult to accomplish the desired whole-genome sequencing of every object in a large cohort. In this scenario, imputation-based strategies, which impute low-density panels to higher densities, offer an alternative to systematic genotyping or sequencing [9, 10]. To date, array-based genotype imputation has been widely used in agricultural species [11, 12]. The imputation accuracy of this strategy crucially depends on the reference panel sizes and genetic distances from the target population. However, the unavailability of large reference panels and array designs for target populations in agricultural species limits the improvement of array-based genotype imputation [13, 14]. Inaccurate imputations influence the results of follow-up analyses, such as genome-wide association studies (GWAS) and genomic selection (GS).

In the recently-developed methods, low-coverage sequencing (LCS) of a large cohort has been proposed to be more informative than sequencing fewer individuals at high coverage [15-17]. Sample sizes and haplotype diversity could be more critical than sequencing depths in determining genotype accuracy of most segregating sites and in increasing the power of association studies. Overall, LCS has been proven to have higher power for trait mapping compared to the array-based genotyping method in human [18]. To date, LCS-based genotype imputation has been employed in many studies using various populations and genotyping algorithms [19-23]. Especially, the STITCH imputation algorithm overcomes the barrier of the lack of good reference panels in non-human species and is applicable even in studies with extremely low sequencing depths [19, 24]. This is a promising approach for agricultural animals without large reference panels in the areas of functional genetic mapping and genomic breeding, but there is no such report yet.

In this study, we describe a cost- and time-efficient low-coverage sequencing method to obtain high-density SNP markers in a large Duroc population [25]. We used the LCS data to demonstrate genotyping and imputation can be inferred with high accuracy in nucleus herds using the BaseVar/STITCH method, allowing further genome-wide association and fine-mapping analyses on multiple traits with high resolution. The LCS strategy provides a powerful way for further exploration of functional genes in agricultural animal breeding.

## Results

### Samples and phenotypes

We choose 2,885 Duroc boars provided by Guangdong Wen’s Foodstuff Group (Guangdong, China) as the study subjects, which were the same samples in the study by Tan et al. [25], and all the pigs were managed on a single nucleus farm. We obtained measurements of four phenotypes with different heritabilities, including back fat thickness at 100 kg (BF), loin muscle area at 100 kg (LMA), lean meat percentage at 100 kg (LMP), and teat number (TN). The estimates of genomic narrow-sense heritability were 0.37±0.05, 0.41±0.05, 0.42±0.06 and 0.37±0.05, respectively. The phenotypic values followed a near bell-shaped distribution (Figure S1 and Table S1).

### Genome sequencing and data acquisition

A Tn5-based protocol was used to prepare sequencing libraries of each pig at low cost (reagent cost: 2.60 $/library) as described in the Materials and Methods. At the beginning, the libraries were sequenced on an Illumina (PE 150) (192 libraries on 2 lanes) or a BGI platform (PE 100) (84 libraries on one lane), and the sequencing depths were 0.40±0.05×/pig for one lane and 0.45±0.06×/pig for the other lane on the Illumina platform and 0.66±0.16×/pig on the BGI platform. The results generated by the BGI platform had lower PCR duplicates (2.23%), higher good index reads (97.10%), and higher genome coverage (98.55%) than the Illumina dataset, which had 10.82% of PCR duplicates, 93.64% of good index reads, and 98.50% of genome coverage. The high PCR duplicates would cause a greater number of useless data, leading to a lower depth for each individual pig. Therefore, the remaining samples were all sequenced on the BGI MGISEQ-2000 platform (96 samples/2 lanes). Overall, the total output of the 2869 boars approached 5.32 TB, and the majority (96.74%) of reads were successfully mapped to the pig reference genome Sscrofa11.1. Each animal was sequenced at an average of depth of 0.73±0.17×, and all the samples had lower levels of PCR duplicates on the BGI platform (2.60±0.08%). Moreover, we also re-sequenced 37 Duroc boars (the core boars of this population) at a high depth (average 10×/per sample), which were used for downstream accuracy evaluation.

### Processing pipeline of the low-coverage strategy and accuracy evaluation

Previous standard methods for joint SNP calling, such as those implemented in GATK and Samtools, were mainly used in high-depth resequencing methods. However, due to the low depth of each base, erroneous SNPs and genotypes could be called using such methods, especially for the GATK HaplotypeCaller algorithm [26]. In this study, we applied the BaseVar algorithm [27] to call SNP variants and estimate allele frequencies, and used STITCH [19] to impute SNPs. The initial screening of chromosome 18 (SSC18) in 1985 samples with BaseVar identified 506,452 and 414,160 bi-allelic candidate SNP sites before and after quality control, respectively. Next, we imputed these SNPs using STITCH, and 322,386 SNPs were retained with a high average call rate (98.89%±0.59%) after quality control. Meanwhile, we also used the GATK UnifiedGenotypeCaller algorithm (different from GATK HaplotypeCaller algorithm) and Beagle to analyze the data and compared the two results (Figure 1). The SNPs detected by BaseVar/STITCH were mostly included (99.32%) in the GATK set, which included 570,919 sites and contained 320,199 SNPs overlapping with the BaseVar/STITCH dataset. To evaluate imputation accuracy, we compared the genotypic concordance (GC) and the allele dosages R^2^ [28] between the genotypes called in the high coverage whole-genome sequencing analyses of the 37 core boars (high-coverage set) and the imputed genotypes in the low-coverage data (LC set). As a result, a relative high-quality genotype set was acquired with less time consumption when K=10 (Figure S2). We then compared the results generated from the GATK-Beagle and BaseVar-STITCH pipelines in parallel. Figure 2 shows that highly accurate genotypes were obtained using the BaseVar-STITCH pipeline (R ^2^=0.92 and GC=0.97) across all allele frequencies, which excelled far beyond the method using GATK and Beagle (R^2^=0.48 and GC=0.71). Therefore, we conclude the BaseVar-STITCH pipeline is a suitable variant discovery and imputation method for the LCS strategy (Figure 1).

**Figure 1.**
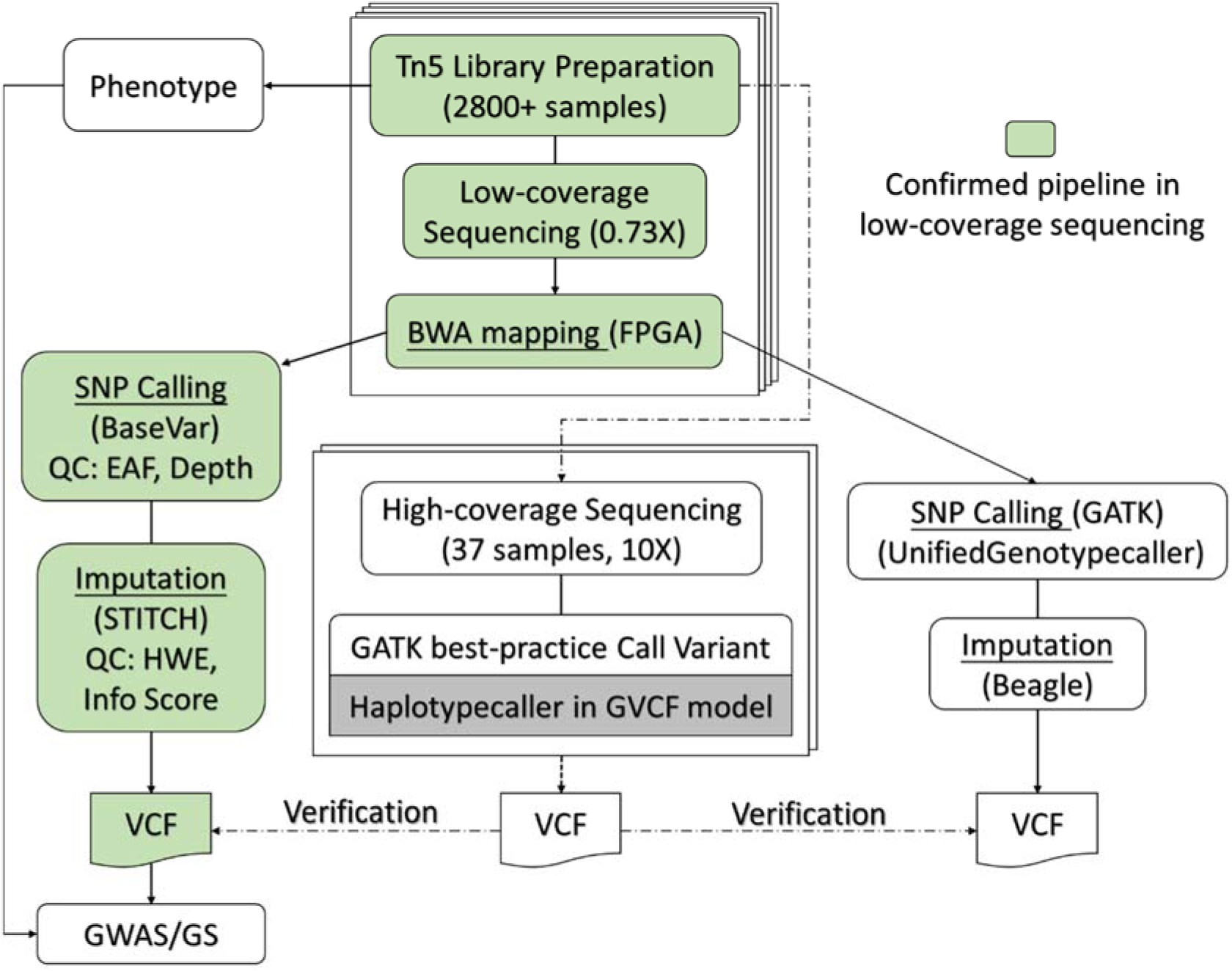
The LCS study design. The flow chart summarizes the steps used to detect and impute SNPs, where the green block represents the pipeline for the LCS analysis (BaseVar-STITCH). The data generated from the GATK-Beagle pipeline were compared with that of the BaseVar-STITCH pipeline, and the data generated from the high-coverage sequencing analyses were used to verify the above results. The BaseVar-STITCH pipeline was used in the further GWAS study.

**Figure 2.**
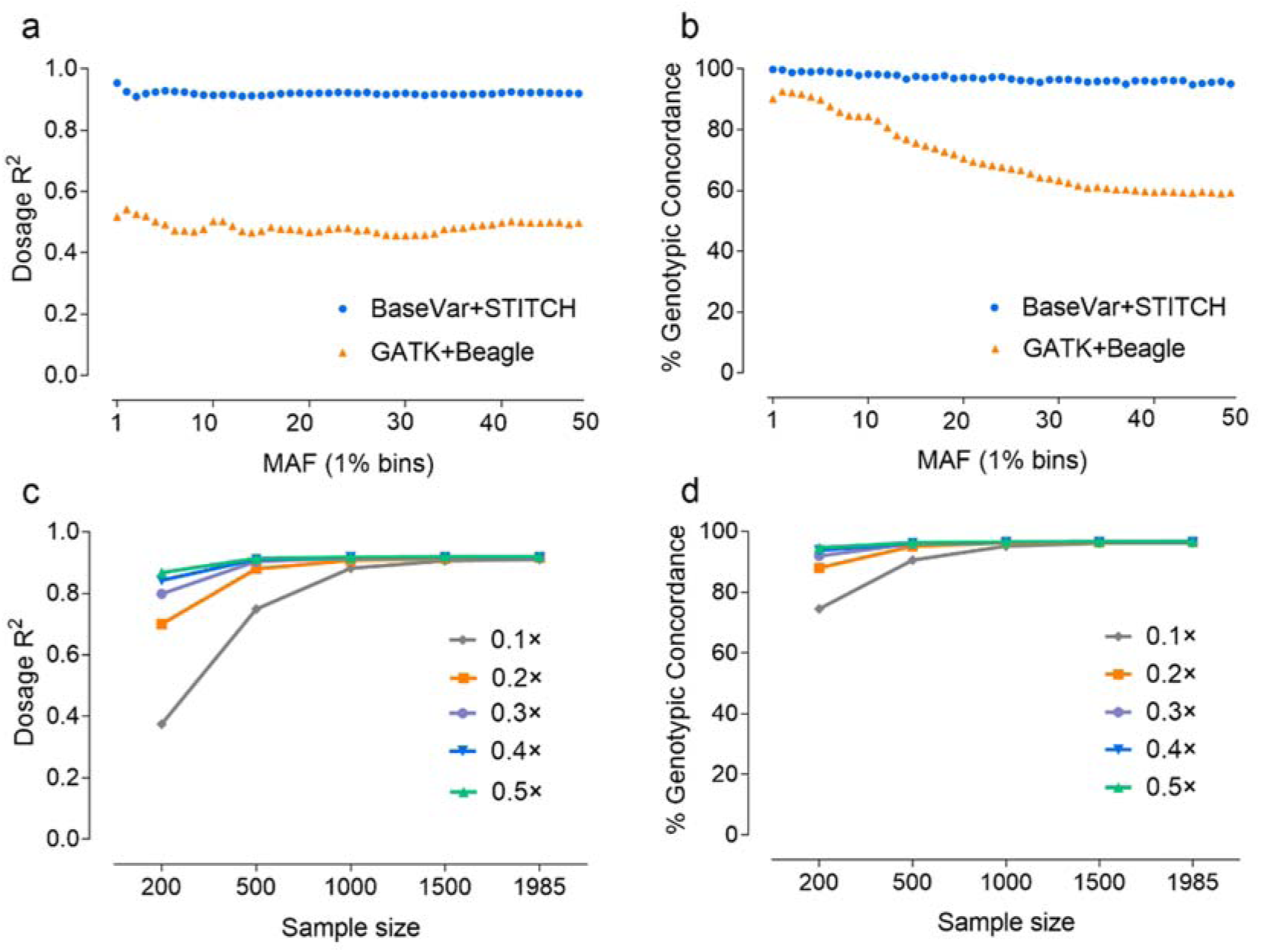
Dosage R^2^ and genotypic concordance (%) values for different MAFs and sample sizes. (a) and (b) show the comparison of Dosage R^2^ and genotypic concordance values between the BaseVar/STITCH for LGS (blue) and the GTAK/Beagle (orange) pipelines, and (c) and (d) show the comparison of Dosage R^2^ and genotypic concordance values among different sequencing depths.

Previous studies have demonstrated that sequencing a large number samples at a low depth generally provides a better representation of population genetic variations compared to sequencing a limited number of individuals at a higher depth. Here, using the proper detection and imputation techniques, we obtained SNP sets based on different depths and sample sizes. From Figure 2, we can see that almost all variants had high concordance and r^2^ values (R^2^>0.91 and GC>0.96) at all depths when the sample size reached 1000. Even at 0.2× sequencing depth, the SNPs were still detected and imputed with high confidence. Therefore, we conclude that both common and low-frequency SNPs (MAF>0.01) can be obtained with high confidence using information from a larger population in the LC strategy, even when the sequencing depth is around 0.2-0.3×.

### Genetic variations and population structure

After strict parameter filtering in the pipeline (BaseVar-STITCH, Figure 1), we identified 11,348,460 SNPs in 2885 Duroc pigs with high genotype accuracy (R^2^=0.92 and GC=0.97), and the density is corresponding to 1 SNP per 200 bp in the pig genome (Table S2). The distribution of variants across the whole genome is mostly uniform, which reflects the high robustness of the LC method. Among all the discovered SNPs, 1,524,015 (accounting for 13.43% of all SNPs) are novel to the pig dbSNP database (data from NCBI:GCA_000003025.6 on Jun, 2017). The majority of identified SNPs were located in intergenic regions (51.98%) and intronic regions (36.85%). The exonic regions contained 1.37% of SNPs, including 0.14% missense SNPs.

Interrogating the distribution of the 11.35 million variants in this Duroc population revealed several genetic characteristics. A principal component analysis (PCA) of all the pigs manifests that there was no distinct population stratification among the population (Figure S3). The genome-wide allele frequency spectrum was shown in Figure 3a. The average rate of heterozygosity is low, which is 0.31 of the genomes (Figure 3b), suggesting a strong selection for the pure-bred population. Based on the large population with high-density SNPs, we analyzed the LD decay. The result showed the average pairwise LD r^2^ decreased slowly along with the increase of distance between markers. The average r^2^ of the whole genome had been decreased to 0.14 when the distance reached 1 Mb, and slight differences in the average r^2^ existed among 18 chromosomes (Figure 3c and Table S3). Overall, using such high-quality and high-density variants, we could obtain more powerful results from GWAS analyses.

**Figure 3.**
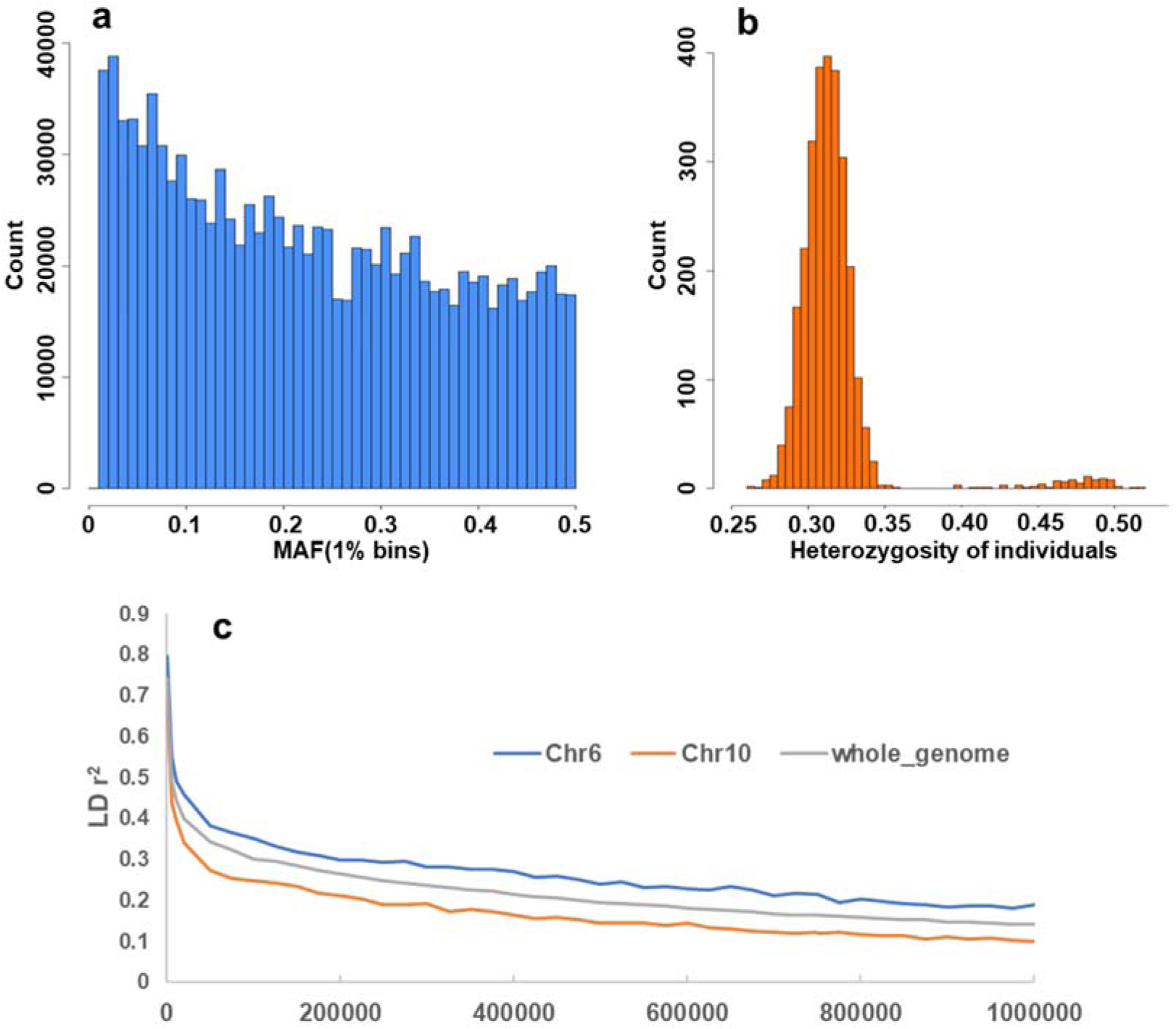
Sequencing diversity of the Duroc population. (a) Histogram of allele counts by each 1% MAF bin. (b) Histogram of genome-wide heterozygosity. (c) The extent of linkage disequilibrium (LD), in which the LD on chromosome 6 and 10 represent the highest and lowest levels among the whole genome respectively.

### Identification of candidate genes by high-resolution mapping QTLs for TN and BF

We identified a subset of 258,662 SNPs that tagged all other SNPs with MAF >0.1% at LD r^2^ >0.98 for the first-round GWAS (Table S2). Fine-mapping was performed within 10 Mb of the SNPs to reach 5% Bonferroni-corrected significance threshold genome-wide. Overall, we discovered a total of seven QTLs for the four traits at 5% significance threshold (Figure 4 and Figure S4). The widths of all QTLs’ intervals ranged from 40 Kb to 3 Mb; the intervals of five QTLs were more than 2 Mb in width, which was strongly influenced by the local linkage disequilibrium level of this population. In the subsequent analyses, we focused on QTLs containing small numbers of genes (TN and BF’s QTLs on SSC7, Figure 4a and 4b).

**Figure 4.**
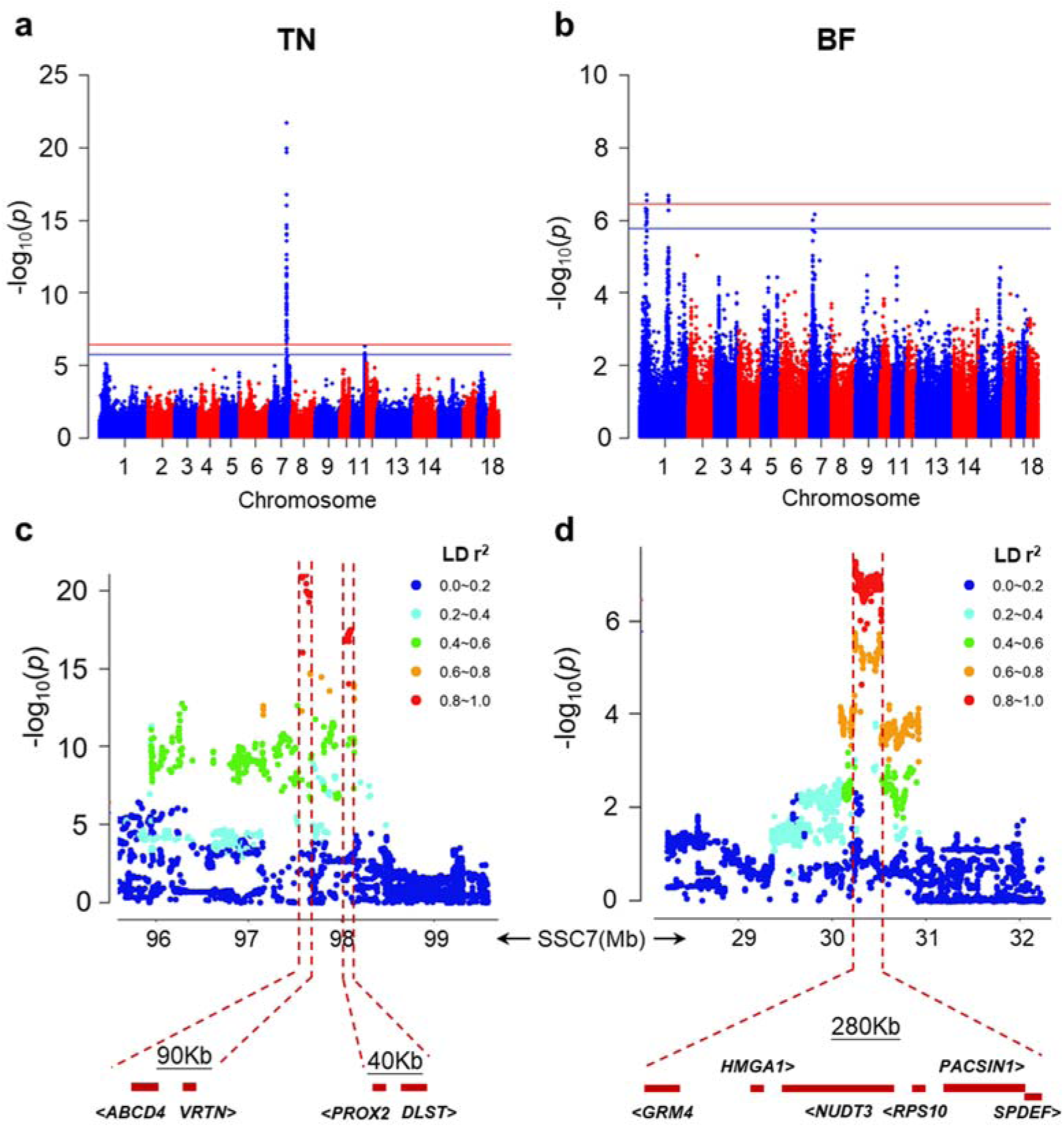
Manhattan plots and fine-mapping of total tit number (TN) and back fat thickness (BF). (a) and (b) depict the TN and BF association signals on the whole genome, in which the blue and red horizontal line represent the 0.05 (p < 1.73×10^−6^) and 0.01 (p < 3.46×10^−7^) significant levels after Bonferroni correction. (c) Fine-mapping of TN using the entire set of SNPs, in which two isolated regions on chromosome 7 with the lengths of 90 Kb and 40 Kb were detected as QTLs. (d) Fine-mapping of BF using the entire set of SNPs, a narrow QTL with the length of 280 Kb on chromosome 7 was detected.

The QTL on SSC7 that has major effect on TN has been widely identified in several commercial breeding lines and hybrids. Our GWAS results show a strong QTL in the same region (Figure 4a). Fine-mapping discovered two narrow LD blocks (SSC7:97.56-97.65 Mb and 98.06-98.10 Mb) containing four candidate genes (Figure 4c). Comparing with the previous results based on the GBS (genotyping-by-sequencing) method, we directly detected 7 missense variants in three genes (*ABCD4, PROX2*, and *DLST*). Besides, our result identified the locus of SSC7: 30.24-30.52 Mb was significantly associated with BF. No missense mutation has been detected in this region, but some UTR variants within six genes may have great effects on this trait. All of these genes, *GRM4, HMGA1, NUDT3, RPS10, PACSIN1*, and *SPDEF* have been reported to be associated with one or multiple traits in pigs, but clear causal mutations still lack. Our results provide a starting point for further functional investigations.

## Discussion

To our knowledge, we generated the largest WGS genotyping data set of Duroc population so far, which contains 11 million markers from genotyping 2885 pigs. We expanded the candidate causal mutations for the TN and BF growth-related traits of pigs and demonstrated the efficacy of genetic fine-mapping utilizing low-coverage sequencing in animal populations with unavailable reference panels. This method is expected to have widespread usages in genome-wide association studies, fine mapping of QTLs, and genomic selection.

This study identified an optimal design, taking into account the number of samples, sequencing depth, and imputation algorithm. Two critical data can be referenced for future research on animals without large reference panels: the BaseVar-STITCH pipeline allows the GC higher than 0.96 when the sample size of 1000 and the sequencing depth of 0.2× were reached, or when the sample size of 500 and the sequencing depth of 0.5× were reached. The GC values under both conditions are significantly higher than other studies of array-based genotype imputation. We also found that the genotype accuracy is more sensitive to sample sizes than sequencing depths. In other words, the results demonstrated low-coverage designs are more powerful than deep sequencing of fewer individuals for animal sequencing studies.

The QTL region on SSC7, which was identified in the current study, has also been reported to be associated with TN, the number of vertebrae (NVE), or the number of ribs by GWASs [25, 29-31]. Vertebrae develop from the somites, whose ventral elongation also determines the correct dorsoventral position of mammary epithelium along the flank [32]. Thus, somites may be the progenitor cells of vertebrae, ribs, and mammary glands, and the variations in the genes downstream of the developmental cascade for the formation of the mammary gland are most likely responsible for the QTL we detected for TN in the GWAS [33]. It is worth noting that five missense variants are discovered in *PROX2*, which is one of the vertebrate homologs of Drosophila melanogaster homeodomain-containing protein Prospero, and may be involved in the determination of cell fate and the establishment of the body plan [34]. We suggest these missense mutations may be the causal variants for the phenotype, although functional studies are needed to validate this hypothesis. For the QTL associated with BF, we found three UTR variants located in *HMGA1. HMGA1* is a promising candidate gene associated with growth, carcass, organ weights, fat metabolism, as it has been reported to involve in a variety of genetic pathways regulating cell growth and differentiation, glucose uptake, and white and brown adipogenesis [35-39]. Overall, the QTLs with narrow intervals and a few candidate genes were identified, which emphasizes the potential of identifying new mutations in QTLs using the low-coverage sequencing method in a single sequencing step.

Increasing the marker density was proposed to have the potential to improve the accuracy of genomic prediction for quantitative traits [40]. However, in recent studies, SNP chips were mostly used to build genetic relationship matrices [41, 42], which could not catch all recombination events in a given population. Here, the whole-genome low-coverage sequencing data gave the best accuracy of prediction, since most causal mutations that underlie a trait are expected to be included. Meanwhile, the haplotype reference panel can accommodate new haplotypes due to recombination at any time, thus improving the issue of the decrease of prediction accuracy over generations. Our data can cover the sites of various of SNP chips well because the genome coverage exceeds 98.36%, and it is competitive with arrays in terms of cost and SNP density. Besides, most researchers or breeders may concern more about the efficiency of the method. The development of application servers brings hope to solve the time-consuming computational issue of genotyping using the whole genome sequencing data. In this study, we applied GTX, which is an FPGA-based hardware accelerator platform [43], to do the alignments, and all 3000 alignments were accomplished in two days. Then the genotyping and imputation could be achieved on the cluster server or even cloud server in a single day. Therefore, the accuracy and timeliness issue for genomic prediction could be all resolved in the near future. An alternative solution at present is that we can select different useful tag-SNPs to make ultra-low-density SNP chips for various traits with different genetic architectures using the high-density genetic map built by LC data, since all possible haplotypes were available in the haplotype database. Further, the cost could be reduced, and breeding could be achieved more efficiently.

## Methods

### Ethics Statement

All procedures involving animals in this study were carried out in accordance with the guidelines for the care and use of experimental animals established by the Ministry of Science and Technology of the People’s Republic of China (Approval Number: 2006-398). All the animal experiment protocols were approved by the Animal Welfare Committee of China Agricultural University (Permission Number: SKLAB-2014-04-02).

### Animals, phenotyping, and DNA Extraction

The Duroc boars used for this study were provided by Guangdong Wen’s Foodstuff Group (Guangdong, China), which were born from September 2011 to September 2013. All pigs were managed on a single nucleus farm. The associated phenotype data used in this study included teat number (TN), back fat thickness at 100 kg (BF), loin muscle area at 100 kg (LMA), and lean meat percentage at 100 kg (LMP). The last three phenotypes were recorded when the weights of pigs reached 100 ± 5 Kg. The phenotype data of TN were acquired from Tan’s study, and BF, LMA, and ELMP were measured over the last three to four ribs using a b-ultrasound-scan equipment (Aloka SSD-500). The phenotypic values of TN followed a near bell-shaped distribution, which is same as reported by Tan et al., and the data of other three phenotypes all nearly followed the normal distribution (Fig S1). In addition, body weights were recorded at birth, and at the beginning (30 ± 5 Kg) and the end (100 ± 5 Kg) of the experiment. Genomic DNA was extracted from the ear tissue using a DNeasy Blood & Tissue Kit (Qiagen 69506), assessed using a NanoDrop, and checked in a 1% agarose gel. All the samples were quantified using a Qubit 2.0 Fluorometer, and then diluted to 40 ng/ml in 96-well plates.

### Tn5 Libraries generation and sequencing

Equal amounts of Tn5ME-A/Tn5MErev and Tn5ME-B/Tn5MErev were incubated at 72 °C for 2 min, then were placed on ice immediately. Tn5 (Karolinska Institutet 171 77 Stockholm, Sweden) was loaded with the Tn5ME-A+rev and Tn5ME-B+rev in 2× Tn5 dialysis buffer at 25°C for 2 h. All linker oligonucleotides were same as the previous report [44].

Tagmentation were carried out at 55°C for 10 minutes by mixing 4 μl 5×TAPS-MgCl_2_, 2 μl dimethylformamide (DMF) (Sigma Aldrich), 1 μl of the Tn5 that pre-diluted to 16.5 ng/μl, 50 ng DNA, and nuclease-free water. The total volume of the reaction was 20 μl. Then 3.5 μl 0.2% SDS was added, and Tn5 was inactivated for another 10 min at 55°C.

KAPA HiFi HotStart ReadyMix (Roche) was used for PCR amplification. The primers were designed for MGI sequencers, with the reverse primers contained 96 different index adaptors to distinguish individual library. The PCR program was as follows: 9 min at 72°C, 30 sec at 98°C, and then 9 cycles of 30 sec at 98°C, 30 sec at 63°C, followed by 3 min at 72°C. The products were quantified by Qubit Fluorometric Quantitation (Invitrogen), then the groups of 96 indexed samples were pooled with equal amounts.

Size-selection was performed using the AMPure XP beads (Beckmann), with the left side size selection ratio was 0.55×, and the right was 0.1×. The final libraries were sequenced on 2 lanes of MGISEQ-2000 to generate 2×100bp paired-end reads or on 1 lane of BGISEQ-500 to generate 2×100 bp paired-end reads.

### High depth sequencing of 37 boars

We sequenced 37 out of the total 1985 pigs using the Hiseq X Ten system at a high depth of 10×. GTX by Genetalks company, a commercially available FPGA-based hardware accelerator platform, was used in this study for both mapping clean reads to the Sscrofa11.1 reference genome (ftp://ftp.ensembl.org/pub/release-97/fasta/sus_scrofa/dna/) and variant calling. The alignment process is accelerated by FPGA implementation of a parallel seed-and-extend approach based on the Smith-Waterman algorithm, while the variant calling process is accelerated by FPGA implementation of GATK HaplotypeCaller (PairHMM). GATK multi-sample best practice [45] was used to call and genotype SNPs for the 37 pigs, and the SNPs were hard filtered with a relatively strict option “QD < 10.0 ‖ ReadPosRankSum < −8.0 ‖ FS > 10.0 ‖ MQ<40.0”. The average running time from a fastq file to a gvcf file was about 3 min for each sample in this study.

### Low coverage sequencing data analyses

Sequencing reads from the low coverage samples were mapped to Sscrofa11.1 reference genome using GTX-align, which includes a step of marking PCR duplicates. The indel realignment and base quality recalibration modules in GATK were applied to realign the reads around indel candidate loci and to recalibrate the base quality. Variant calling was done using the BaseVar and hard filtered with EAF >= 0.01 and the Depth that is greater than or equal to 1.5 times InterQuartile Range. The detailed BaseVar algorithm to call SNP variants and to estimate allele frequency was described in a pevious report [27]. We used STITCH [19] to impute genotype probabilities for all individuals. The key parameter K (number of ancestral haplotypes) was decided based on the tests in SSC18. Results were filtered with an imputation info score > 0.4 and Hardy-Weinberg Equilibrium (HWE) p-value > 1e−6. Two validation actions were taken to calculate the accuracy of imputation. The first parameter is genotypic concordance (GC), which was calculated as the number of correctly-imputed genotypes divided by total sites. Another parameter is allele dosages R^2^, which was described in a previous report [28]. The SNPEff program [46] was used to annotate the variants.

### Heritability estimation and Genome-wide association

Heritability was estimated using a mixed model as the following: 

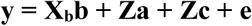

with Var(**y**) = **ZA**_a_**Z’**ơ_a_^2^ **+ I**ơ_c_^2^ **+ I**ơ_e_^2^, where Z is an incidence matrix allocating phenotypic observations to each animal; **b** is the vector of fixed year-month effects for BF, LMA, and ELMP; **b** also includes birth weight, the weights at the beginning and end of the test as covariance; **Xb** is the incidence matrix for **b**; **a** is the vector of additive values based on the pedigree information; **c** is the vector of random family effects; **A**_a_ is a pedigree-based additive relationship matrix; ơ_a_^2^ is the additive variance; ơ_c_^2^ is the variance of random family effects; and ơ_e_^2^ is the residual variance. Variance components for BF, LMA, and LMP were estimated by AIREMlF90 program, and by thrgibbs1f90 program for TN. Both programs were in the BLUPF90 package. The additive heritability was defined as: h_a_^2^ = ơ_a_^2^ /(ơ_a_^2^ + ơ_c_^2^ + ơ_e_^2^).

A subset of 258,662 SNPs that tagged all other SNPs with MAF >1% at LD r^2^ <0.98 and the call rate >95% were retained for genome-wide association analysis. A mixed linear model (MLM) approach was used for the genome-wide association analyses as implemented in the GCTA package (v1.24) [47]. The statistical model during analyses of TN included the year and season as discrete covariates. For BF, LMA, and ELMP, the year and season were included as discrete covariates, and birth weight, the weight at the beginning and end of the test were used as quantitative covariate. To correct multiple testing across the genome, a Bonferroni correction was applied to compensate for the number of estimated independent markers from a PCA analysis and was performed as follows. A subset of SNPs that were in approximate linkage equilibrium with each other was obtained by removing one in each pair of SNPs if the LD was greater than 0.5 using the PLINK v1.07 ‘--indep-pairwise’ command [48]. The squared correlation coefficient (r^2^) between the genotypes was calculated using the vcftools ‘--geno-r2’ command [45]. Consequently, for our population, the genome-wide 1% significance threshold was determined as p-value < 3.47×10^−7^ (0.01/28,828), and a suggestive association was determined as 1.73 ×10^−6^ (0.05/28,828).

## Acknowledgements

We thank Siyang Liu and Xun Xu for their valuable suggestions on data analyses. Zhuo Song, Chungen Yi, and Wenjuan Wei provided the services of FPGA-based hardware accelerator platform and the Batch Compute system in Aliyun cloud. We also thank Zhaoliang Liu for improving the manuscript. Part of the analysis was performed on the high-performance computing platform of the State Key Laboratory of Agrobiotechnology.

