## Supplementary for "Genome-wide association analyses of multiple traits in Duroc pigs using low-coverage whole-genome sequencing strategy"

**Table S1 Summary of four traits**

| | BF | TN | LMA | LMP |
|---|---|---|---|---|
| Individuals | 2800 | 2803 | 2801 | 2801 |
| Heritability | 0.373±0.057 | 0.377±0.045 | 0.414±0.056 | 0.418±0.059 |
| Phenotypes (Min/Max) | 6.10/20.73 (mm) | 8.00/15.00 | 21.75/51.04 ($mm^2$) | 47.15/59.89 |
| Phenotypes (MEAN±STD) | 11.22±2.07 (mm) | 10.73±1.07 | 36.24±3.61 ($mm^2$) | 54.02±1.58 |

**Table S2 Number and density of SNPs imputed by STITCH and Tag SNPs used in GWAS**

| Chr. | Size | Clean SNP of STITCH | Density (N/Mb) | Tag SNP for GWAS | Density (N/Mb) |
|---|---|---|---|---|---|
| 1 | 274330532 | 884568 | 3224.46 | 19264 | 70.22 |
| 2 | 151935994 | 730256 | 4806.34 | 15751 | 103.67 |
| 3 | 132848913 | 652638 | 4912.63 | 15692 | 118.12 |
| 4 | 130910915 | 698766 | 5337.72 | 17731 | 135.44 |
| 5 | 104526007 | 536643 | 5134.06 | 13997 | 133.91 |
| 6 | 170843587 | 705146 | 4127.44 | 14780 | 86.51 |
| 7 | 121844099 | 603195 | 4950.55 | 13374 | 109.76 |
| 8 | 138966237 | 871714 | 6272.85 | 17421 | 125.36 |
| 9 | 139512083 | 674562 | 4835.15 | 15446 | 110.71 |
| 10 | 69359453 | 578249 | 8336.99 | 14756 | 212.75 |
| 11 | 79169978 | 524580 | 6626.00 | 13316 | 168.20 |
| 12 | 61602749 | 422758 | 6862.65 | 9366 | 152.04 |
| 13 | 208334590 | 796182 | 3821.65 | 16822 | 80.75 |
| 14 | 141755446 | 752846 | 5310.88 | 16443 | 116.00 |
| 15 | 140412725 | 676856 | 4820.47 | 14288 | 101.76 |
| 16 | 79944280 | 483771 | 6051.35 | 12077 | 151.07 |
| 17 | 63494081 | 450491 | 7095.01 | 11943 | 188.10 |
| 18 | 55982971 | 305239 | 5452.35 | 6195 | 110.66 |
| Total | 2265774640 | 11348460 | 5008.64 | 258662 | 114.16 |

**Table S3 The average pairwise LD $r^2$ among 18 chromosomes**

| distance/bp | Chr1 | Chr2 | Chr3 | Chr4 | Chr5 | Chr6 | Chr7 | Chr8 | Chr9 | Chr10 | Chr11 | Chr12 | Chr13 | Chr14 | Chr15 | Chr16 | Chr17 | Chr18 |
|---|---|---|---|---|---|---|---|---|---|---|---|---|---|---|---|---|---|---|
| 10 | 0.78 | 0.70 | 0.75 | 0.71 | 0.75 | 0.80 | 0.80 | 0.72 | 0.76 | 0.71 | 0.72 | 0.74 | 0.75 | 0.73 | 0.74 | 0.70 | 0.71 | 0.79 |
| 5000 | 0.54 | 0.44 | 0.50 | 0.44 | 0.49 | 0.55 | 0.55 | 0.48 | 0.51 | 0.44 | 0.47 | 0.48 | 0.51 | 0.47 | 0.49 | 0.45 | 0.44 | 0.60 |
| 10000 | 0.49 | 0.40 | 0.47 | 0.39 | 0.43 | 0.49 | 0.49 | 0.43 | 0.46 | 0.40 | 0.41 | 0.45 | 0.47 | 0.43 | 0.44 | 0.39 | 0.41 | 0.53 |
| 20000 | 0.44 | 0.38 | 0.41 | 0.35 | 0.38 | 0.46 | 0.45 | 0.39 | 0.41 | 0.34 | 0.36 | 0.39 | 0.42 | 0.39 | 0.41 | 0.36 | 0.37 | 0.48 |
| 50000 | 0.38 | 0.35 | 0.35 | 0.30 | 0.33 | 0.38 | 0.41 | 0.33 | 0.36 | 0.27 | 0.31 | 0.35 | 0.38 | 0.34 | 0.34 | 0.30 | 0.32 | 0.39 |
| 75000 | 0.37 | 0.32 | 0.34 | 0.28 | 0.30 | 0.36 | 0.36 | 0.31 | 0.34 | 0.25 | 0.28 | 0.33 | 0.35 | 0.31 | 0.34 | 0.29 | 0.30 | 0.37 |
| 100000 | 0.34 | 0.30 | 0.32 | 0.28 | 0.28 | 0.35 | 0.35 | 0.29 | 0.31 | 0.25 | 0.27 | 0.31 | 0.32 | 0.31 | 0.31 | 0.26 | 0.28 | 0.33 |
| 125000 | 0.34 | 0.30 | 0.31 | 0.27 | 0.28 | 0.33 | 0.34 | 0.28 | 0.32 | 0.24 | 0.25 | 0.29 | 0.32 | 0.29 | 0.30 | 0.26 | 0.28 | 0.31 |
| 150000 | 0.32 | 0.27 | 0.30 | 0.26 | 0.28 | 0.32 | 0.34 | 0.28 | 0.29 | 0.23 | 0.25 | 0.30 | 0.31 | 0.29 | 0.30 | 0.24 | 0.27 | 0.29 |
| 175000 | 0.31 | 0.27 | 0.29 | 0.24 | 0.26 | 0.31 | 0.33 | 0.27 | 0.28 | 0.22 | 0.23 | 0.27 | 0.29 | 0.28 | 0.29 | 0.24 | 0.25 | 0.29 |
| 200000 | 0.31 | 0.29 | 0.28 | 0.23 | 0.25 | 0.30 | 0.31 | 0.25 | 0.28 | 0.21 | 0.24 | 0.27 | 0.29 | 0.26 | 0.28 | 0.23 | 0.24 | 0.27 |
| 225000 | 0.29 | 0.26 | 0.27 | 0.22 | 0.25 | 0.30 | 0.32 | 0.25 | 0.27 | 0.20 | 0.22 | 0.26 | 0.27 | 0.26 | 0.27 | 0.22 | 0.23 | 0.25 |
| 250000 | 0.29 | 0.26 | 0.26 | 0.22 | 0.23 | 0.29 | 0.29 | 0.23 | 0.26 | 0.19 | 0.23 | 0.26 | 0.27 | 0.25 | 0.26 | 0.22 | 0.23 | 0.24 |
| 275000 | 0.28 | 0.24 | 0.24 | 0.22 | 0.24 | 0.29 | 0.28 | 0.24 | 0.25 | 0.19 | 0.22 | 0.24 | 0.27 | 0.25 | 0.25 | 0.20 | 0.22 | 0.23 |
| 300000 | 0.27 | 0.24 | 0.24 | 0.21 | 0.22 | 0.28 | 0.28 | 0.23 | 0.25 | 0.19 | 0.20 | 0.24 | 0.25 | 0.24 | 0.25 | 0.20 | 0.22 | 0.23 |
| 325000 | 0.27 | 0.24 | 0.24 | 0.21 | 0.22 | 0.28 | 0.27 | 0.23 | 0.24 | 0.17 | 0.19 | 0.22 | 0.25 | 0.24 | 0.25 | 0.20 | 0.22 | 0.23 |
| 350000 | 0.26 | 0.24 | 0.22 | 0.20 | 0.21 | 0.28 | 0.27 | 0.22 | 0.23 | 0.18 | 0.20 | 0.22 | 0.25 | 0.24 | 0.24 | 0.19 | 0.21 | 0.20 |
| 375000 | 0.25 | 0.21 | 0.22 | 0.21 | 0.21 | 0.28 | 0.27 | 0.22 | 0.23 | 0.17 | 0.20 | 0.22 | 0.24 | 0.23 | 0.23 | 0.19 | 0.21 | 0.20 |
| 400000 | 0.25 | 0.21 | 0.22 | 0.20 | 0.19 | 0.27 | 0.26 | 0.21 | 0.23 | 0.16 | 0.19 | 0.20 | 0.23 | 0.23 | 0.23 | 0.19 | 0.21 | 0.20 |
| 425000 | 0.24 | 0.22 | 0.21 | 0.20 | 0.20 | 0.26 | 0.25 | 0.21 | 0.23 | 0.16 | 0.18 | 0.20 | 0.22 | 0.22 | 0.23 | 0.17 | 0.19 | 0.18 |
| 450000 | 0.24 | 0.20 | 0.19 | 0.19 | 0.20 | 0.26 | 0.25 | 0.20 | 0.22 | 0.16 | 0.19 | 0.20 | 0.23 | 0.21 | 0.22 | 0.17 | 0.19 | 0.18 |
| 475000 | 0.22 | 0.20 | 0.20 | 0.18 | 0.19 | 0.25 | 0.24 | 0.20 | 0.21 | 0.15 | 0.18 | 0.18 | 0.22 | 0.21 | 0.22 | 0.17 | 0.18 | 0.18 |
| 500000 | 0.22 | 0.20 | 0.19 | 0.18 | 0.19 | 0.24 | 0.24 | 0.20 | 0.21 | 0.14 | 0.18 | 0.18 | 0.22 | 0.20 | 0.22 | 0.17 | 0.17 | 0.18 |

| | | | | | | | | | | | | | | | | | | |
|---|---|---|---|---|---|---|---|---|---|---|---|---|---|---|---|---|---|---|
| 525000 | 0.23 | 0.17 | 0.19 | 0.18 | 0.19 | 0.25 | 0.23 | 0.20 | 0.20 | 0.14 | 0.17 | 0.18 | 0.21 | 0.21 | 0.21 | 0.17 | 0.18 | 0.17 |
| 550000 | 0.22 | 0.17 | 0.19 | 0.17 | 0.18 | 0.23 | 0.24 | 0.19 | 0.20 | 0.15 | 0.17 | 0.17 | 0.20 | 0.19 | 0.20 | 0.17 | 0.18 | 0.16 |
| 575000 | 0.21 | 0.19 | 0.18 | 0.16 | 0.18 | 0.24 | 0.23 | 0.20 | 0.21 | 0.14 | 0.16 | 0.17 | 0.19 | 0.19 | 0.21 | 0.15 | 0.17 | 0.15 |
| 600000 | 0.21 | 0.17 | 0.19 | 0.17 | 0.17 | 0.23 | 0.22 | 0.19 | 0.19 | 0.14 | 0.17 | 0.17 | 0.20 | 0.19 | 0.21 | 0.16 | 0.17 | 0.14 |
| 625000 | 0.21 | 0.17 | 0.17 | 0.16 | 0.17 | 0.23 | 0.22 | 0.18 | 0.20 | 0.13 | 0.15 | 0.16 | 0.19 | 0.19 | 0.21 | 0.15 | 0.16 | 0.14 |
| 650000 | 0.20 | 0.17 | 0.17 | 0.16 | 0.16 | 0.23 | 0.21 | 0.18 | 0.19 | 0.13 | 0.15 | 0.15 | 0.19 | 0.19 | 0.20 | 0.15 | 0.17 | 0.14 |
| 675000 | 0.21 | 0.16 | 0.17 | 0.16 | 0.16 | 0.23 | 0.22 | 0.17 | 0.19 | 0.13 | 0.15 | 0.15 | 0.19 | 0.18 | 0.20 | 0.16 | 0.15 | 0.13 |
| 700000 | 0.19 | 0.15 | 0.17 | 0.16 | 0.15 | 0.21 | 0.21 | 0.18 | 0.18 | 0.12 | 0.15 | 0.15 | 0.18 | 0.18 | 0.20 | 0.14 | 0.15 | 0.14 |
| 725000 | 0.20 | 0.15 | 0.17 | 0.15 | 0.15 | 0.22 | 0.20 | 0.17 | 0.18 | 0.12 | 0.14 | 0.15 | 0.18 | 0.18 | 0.19 | 0.15 | 0.15 | 0.13 |
| 750000 | 0.20 | 0.14 | 0.16 | 0.15 | 0.15 | 0.22 | 0.21 | 0.17 | 0.17 | 0.12 | 0.14 | 0.14 | 0.18 | 0.18 | 0.19 | 0.14 | 0.15 | 0.14 |
| 775000 | 0.20 | 0.15 | 0.17 | 0.15 | 0.15 | 0.20 | 0.21 | 0.17 | 0.18 | 0.12 | 0.15 | 0.14 | 0.18 | 0.17 | 0.19 | 0.14 | 0.14 | 0.13 |
| 800000 | 0.19 | 0.14 | 0.16 | 0.14 | 0.14 | 0.20 | 0.19 | 0.17 | 0.18 | 0.12 | 0.14 | 0.14 | 0.18 | 0.17 | 0.18 | 0.14 | 0.14 | 0.13 |
| 825000 | 0.19 | 0.13 | 0.15 | 0.14 | 0.15 | 0.20 | 0.21 | 0.17 | 0.17 | 0.11 | 0.14 | 0.13 | 0.18 | 0.16 | 0.18 | 0.14 | 0.14 | 0.13 |
| 850000 | 0.18 | 0.13 | 0.16 | 0.14 | 0.14 | 0.19 | 0.20 | 0.16 | 0.17 | 0.11 | 0.13 | 0.14 | 0.17 | 0.16 | 0.18 | 0.13 | 0.13 | 0.12 |
| 875000 | 0.18 | 0.13 | 0.16 | 0.14 | 0.14 | 0.19 | 0.19 | 0.16 | 0.17 | 0.11 | 0.14 | 0.13 | 0.18 | 0.17 | 0.18 | 0.14 | 0.14 | 0.12 |
| 900000 | 0.18 | 0.12 | 0.15 | 0.13 | 0.14 | 0.18 | 0.19 | 0.16 | 0.16 | 0.11 | 0.13 | 0.13 | 0.16 | 0.16 | 0.18 | 0.13 | 0.13 | 0.12 |
| 925000 | 0.18 | 0.13 | 0.15 | 0.14 | 0.14 | 0.19 | 0.19 | 0.15 | 0.16 | 0.11 | 0.13 | 0.12 | 0.16 | 0.15 | 0.17 | 0.12 | 0.14 | 0.12 |
| 950000 | 0.17 | 0.13 | 0.15 | 0.13 | 0.13 | 0.19 | 0.18 | 0.16 | 0.16 | 0.11 | 0.12 | 0.12 | 0.17 | 0.16 | 0.17 | 0.13 | 0.13 | 0.12 |
| 975000 | 0.17 | 0.12 | 0.14 | 0.13 | 0.13 | 0.18 | 0.18 | 0.15 | 0.15 | 0.10 | 0.12 | 0.12 | 0.16 | 0.16 | 0.17 | 0.13 | 0.13 | 0.11 |
| 1000000 | 0.17 | 0.12 | 0.14 | 0.12 | 0.13 | 0.19 | 0.19 | 0.16 | 0.15 | 0.10 | 0.13 | 0.12 | 0.17 | 0.15 | 0.17 | 0.12 | 0.12 | 0.12 |

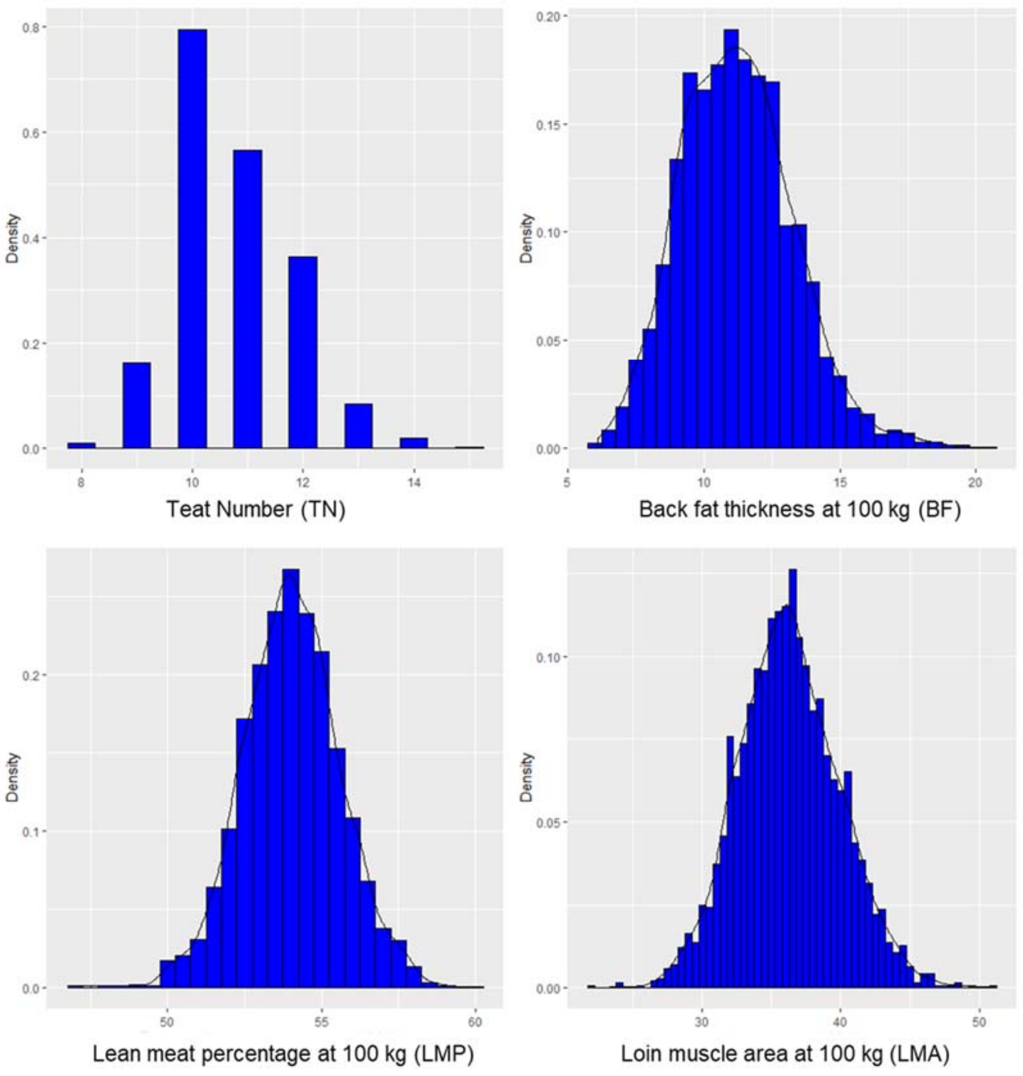


**Figure S1 The data distribution of TN, BF, LMP, and LMA phenotypes**

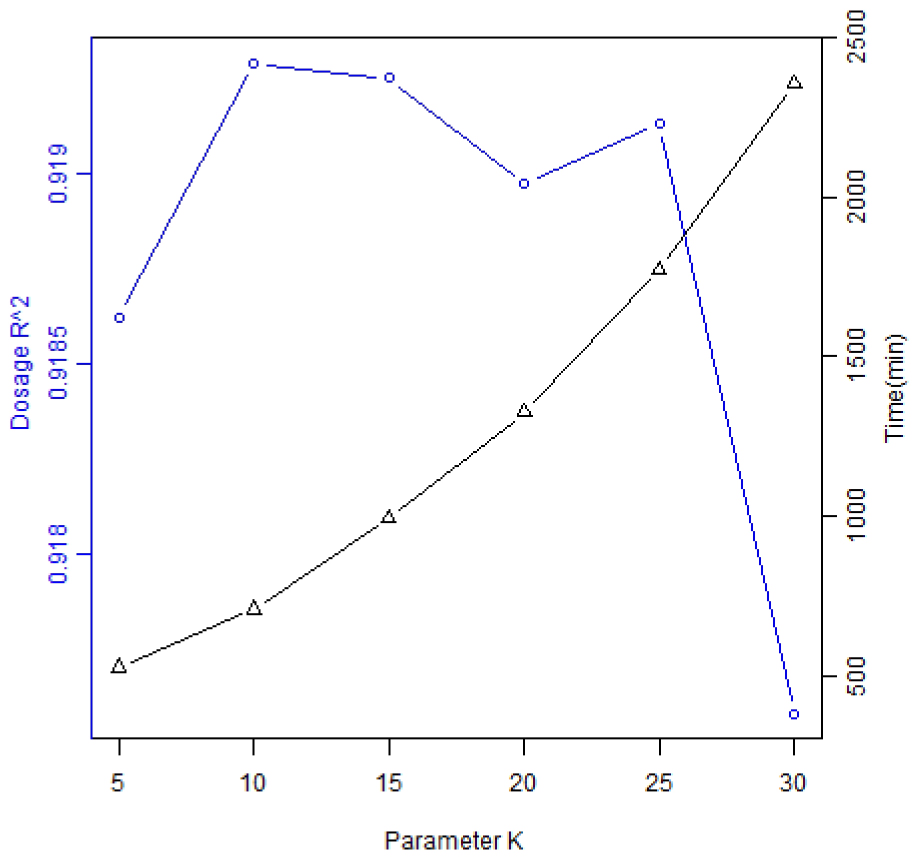


**Figure S2 Dosage $R^2$ and cost time (minute) at different K values. The computation was** tested on chromosome 18, and 20 cores were used for calculation.

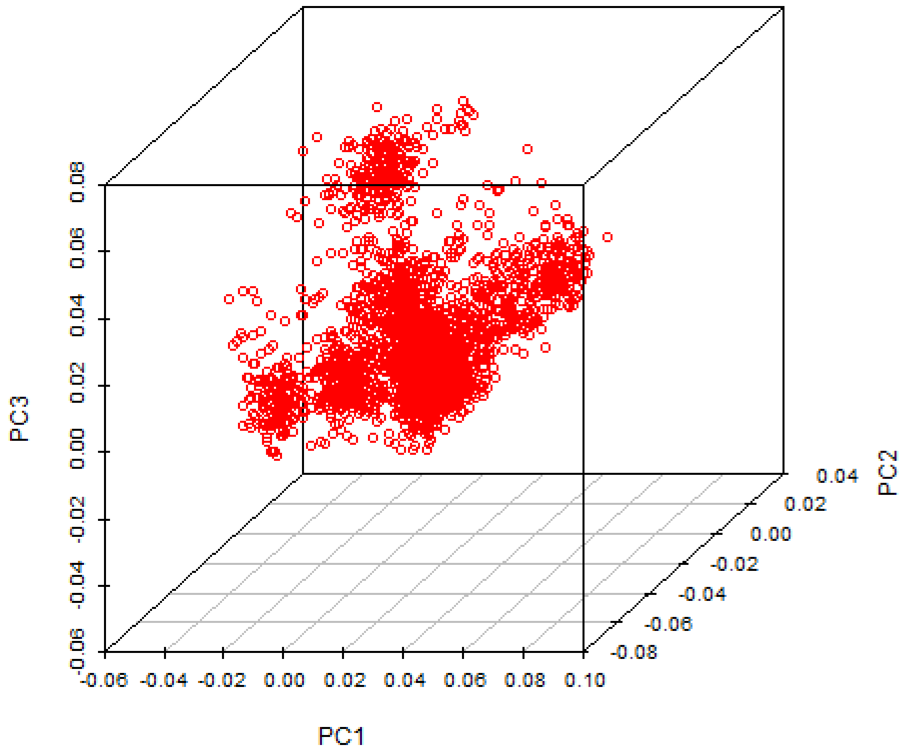


**Figure S3 Principal component 1, 2, and 3 in 2885 pigs**

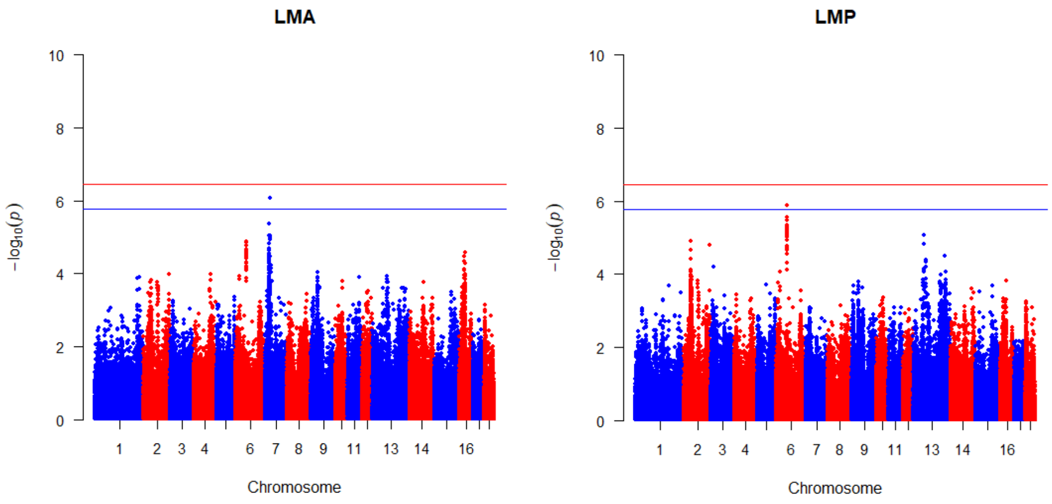


**Figure S4 Manhattan plots of LMA and LMP phenotypes.** The horizontal red/blue lines indicate the genome-wide 1/5% significance thresholds (P values = $3.47 \times 10^{-7}$ and $1.73 \times 10^{-6}$ respectively).
